# The malaria metabolite HMBPP does not trigger erythrocyte terpene production

**DOI:** 10.1101/2020.07.29.225839

**Authors:** Justin J. Miller, Audrey R. Odom John

## Abstract

Infection with malarial parasites renders hosts more mosquito attractive than their uninfected, healthy, counterparts. One volatile organic compound, α-pinene, is associated with *Plasmodium* spp. infection in multiple studies and is a known mosquito attractant. However, how malarial infection results in elevated levels of host-associated α-pinene remains unclear. One study suggests that erythrocyte exposure to the malarial metabolite, (E)-4-hydroxy-3-methyl-but-2-enyl pyrophosphate (HMBPP), results in increased levels of α-pinene. Here, we establish that endogenous levels of α-pinene are present in human erythrocytes, that these levels vary widely by erythrocyte donor, and that α-pinene levels are not altered by HMBPP treatment.

---

*Plasmodium falciparum*, the primary causative agent of lethal malaria infections, has a two-host life cycle between humans and mosquitoes. Transit between the two hosts is a critical requirement for the parasite lifecycle and represents a substantial population bottleneck^1^. Mosquitoes are more attracted to humans^2–5^, mice^6^, and birds^7^ infected with malaria parasites in comparison to uninfected, healthy hosts. This observation has led to the hypothesis that *Plasmodium* species actively manipulate host odor profiles to coordinate transmission to the mosquito. Indeed, changes in the composition of host odor profiles have been observed in humans^8–13^ and mice^6^ infected with malaria; however the molecular basis for infection-induced changes in volatile organic compound (VOC) production or release remains unknown.

Of particular interest has been the mosquito semiochemical, α-pinene, which is found in higher concentrations in the breath of humans with symptomatic *Plasmodium* infection vs healthy controls^8^. Additionally, α-pinene has been identified in the headspace above *Plasmodium falciparum* infected erythrocytes^14^. The VOC α-pinene is a member of the large and bioactive class of molecules termed terpenes. Terpenes are biosynthesized by a variety of plants, soil and environmental organisms, mammalian commensal and pathogenic microbes, and some insects^15–22^. α-pinene is a known component of plant-derived odorant blends that are attractive to the *Anopheles* spp. mosquitoes that transmit malaria^23,24^. As for other terpenes, biosynthesis of α-pinene begins with the 5-carbon isoprenoid precursor, isopentyl pyrophosphate (IPP), which is enzymatically condensed with a second molecule of IPP by geranyl pyrophosphate synthase (GPPS) to form the 10-carbon metabolite, geranyl pyrophosphate (GPP). Subsequent rearrangement and cyclization are catalyzed by a monoterpene synthase (pinene synthase) to yield α-pinene (Fig 1A).

**Figure 1.**
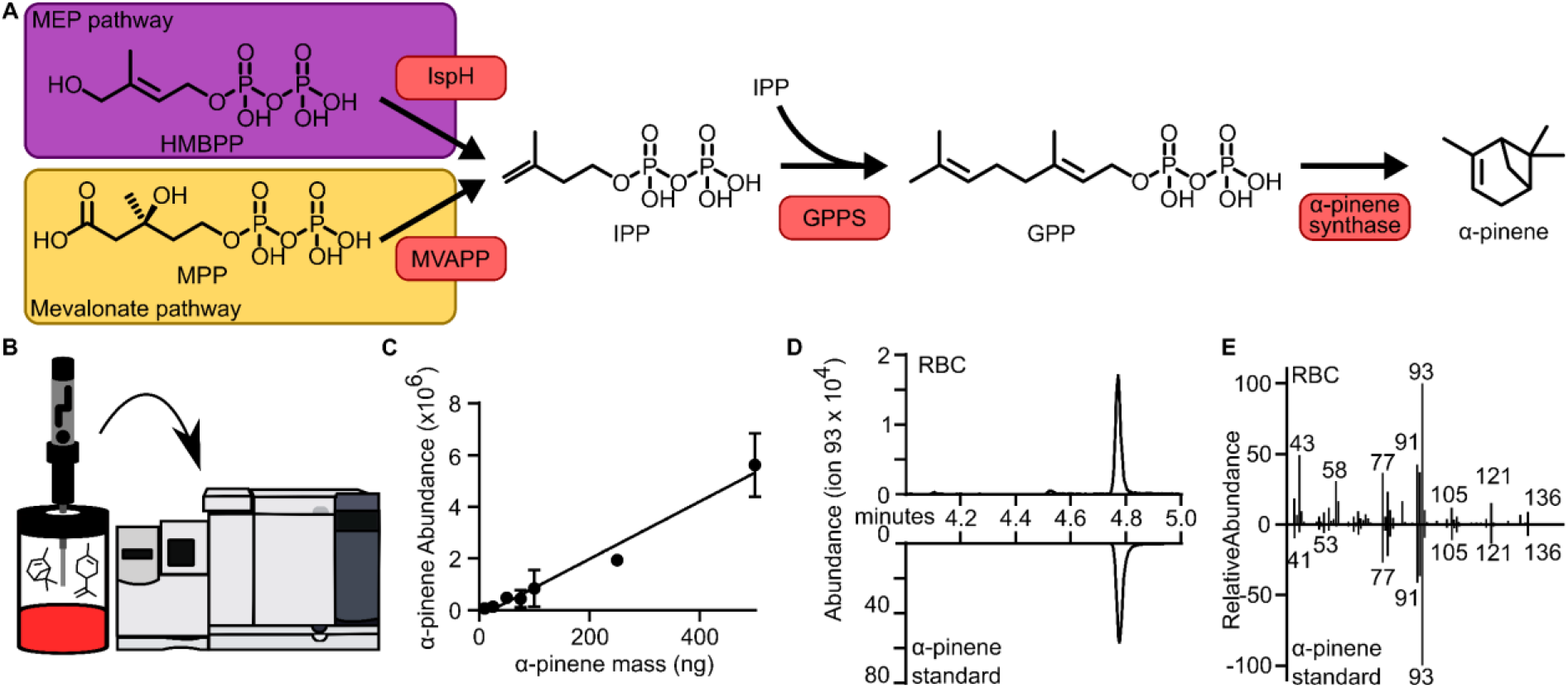
(A) Described metabolic pathways leading to α-pinene. Enzymes highlighted with salmon boxes. (B) Schematic of α-pinene detection assay. (C) α-pinene standard curve generated using commercial α-pinene. (D) gas chromatography-mass spectroscopy trace of commercial α-pinene (bottom) and erythrocyte headspace (top) for α-pinene parent ion, 93. (E) Mass spectra from retention time 4.77 min, the elution time for commercial α-pinene, for erythrocyte headspace (top) and commercial α-pinene (bottom).

Recently, it was reported that incubation of the microbial metabolite (E)-4-hydroxy-3-methyl-but-2-enyl pyrophosphate (HMBPP) with uninfected human erythrocytes results in increased attraction and feeding behavior of anopheline mosquitoes. Concordantly, an increase in the headspace concentration of α-pinene above HMBPP-treated erythrocytes was also reported^25^. HMBPP is a late intermediate in the 2-C-methyl-D-erythritol 4-phosphate (MEP) pathway for synthesis of IPP and downstream isoprenoids (Fig 1A). While eubacteria and apicomplexan parasites, such as *Plasmodium* spp., utilize the MEP pathway for isoprenoid biosynthesis^26^, humans utilize a distinct and evolutionarily divergent biosynthetic pathway (mevalonate pathway) to synthesize IPP^27^.

The mechanism by which HMBPP exposure of erythrocytes may lead to α-pinene production or release is unclear, but two possibilities may explain these findings. First, HMBPP may serve as an exogenous signal that triggers erythrocytes to release stores of α-pinene which may have accumulated via metabolic, environmental, or dietary routes. A potent activator of human Vγ9Vδ2-T cells^28^, HMBPP is recognized as a pathogen-associated molecular pattern (PAMP) through its interaction with butyrophilin receptors^29^, suggesting that HMBPP may serve a signaling role to mediate erythrocyte α-pinene release. Alternatively, because HMBPP is itself a precursor to isoprenoids and terpenes in bacteria and plants, this metabolite may be directly incorporated into α-pinene in erythrocytes via the pathway illustrated in Fig 1A, or via an as-yet-undescribed alternative enzymatic route. However, human erythrocytes do not express the known biosynthetic machinery for synthesis of α-pinene from HMBPP; mammals lack the MEP pathway and specifically do not express the final enzyme in the pathway, IspH, which converts HMBPP to the immediate α-pinene precursor, IPP. Finally, no erythrocyte monoterpene synthases, nor any proteins with the terpene synthase fold, have yet been described that might mediate the final biocatalysis of GPP to α-pinene. In contrast, humans do express other prenyl diphosphate synthase orthologs, and these enzymes have been reported to moonlight as terpene synthases^20–22^. For this reason, we sought to interrogate the possibility of HMBPP-triggered, erythrocyte-produced α-pinene.

We established a working method for sampling the volatile organic compounds associated with cultured erythrocytes. Similar to Emami *et al*., we sealed donated human erythrocytes within a closed, airtight chromatography vial, prewarmed to 38 °C, and performed headspace sampling using solid phase microextraction (SPME) (Fig 1B). Headspace composition was determined using gas chromatography-mass spectrometry. Using a pure commercial α-pinene standard, we established the sensitivity and dynamic range of this assay (Fig 1C), yielding a signal-to-noise ratio of 3 and a limit-of-detection of 0.3 ng α-pinene. Accommodating volumes up to 1 mL of blood, we can detect α-pinene blood concentrations as low as 2.2 nmol/L. We next sought to determine whether α-pinene was present in the headspace above untreated erythrocytes. Indeed, we confirmed that α-pinene can be detected in the headspace from donor erythrocytes, and both the retention time and mass spectra match that of the pure α-pinene standard (Fig 1D,E).

A previous study had indicated that treatment of human erythrocytes with the microbial metabolite HMBPP leads to substantial release of α-pinene. To control for batch-to-batch variability in low-level contaminants present in purified HMBPP, we acquired HMBPP from two independent chemical suppliers. Headspace sampling from both pure preparations of HMBPP confirmed that neither had contaminating levels of α-pinene above our limit-of-detection (Fig 2A). We next treated erythrocytes with either HMBPP or water (vehicle control) and quantified headspace α-pinene. Because monoterpenes such as α-pinene can diffuse into the ambient air, we pre-aliquoted all blood samples into sealed individual-use aliquots. Treatment of erythrocytes with HMBPP did not result in increased levels of α-pinene (Fig 2b), and this finding was not donor-dependent. A previous study also indicated that levels of other monoterpenes (β-pinene and limonene), as well as several aldehydes (octanal, nonanal, and decanal), were increased in response to HMBPP treatment. We evaluated for the presence of additional VOCs including these and found all were below the limit-of-detection for our assay. Our studies thus indicate that if erythrocytes can sense HMBPP, this signal is not accompanied by a substantial release of α-pinene. In addition, our studies suggest that erythrocytes do not incorporate exogenous HMBPP for direct *de novo* synthesis of α-pinene.

**Figure 2.**
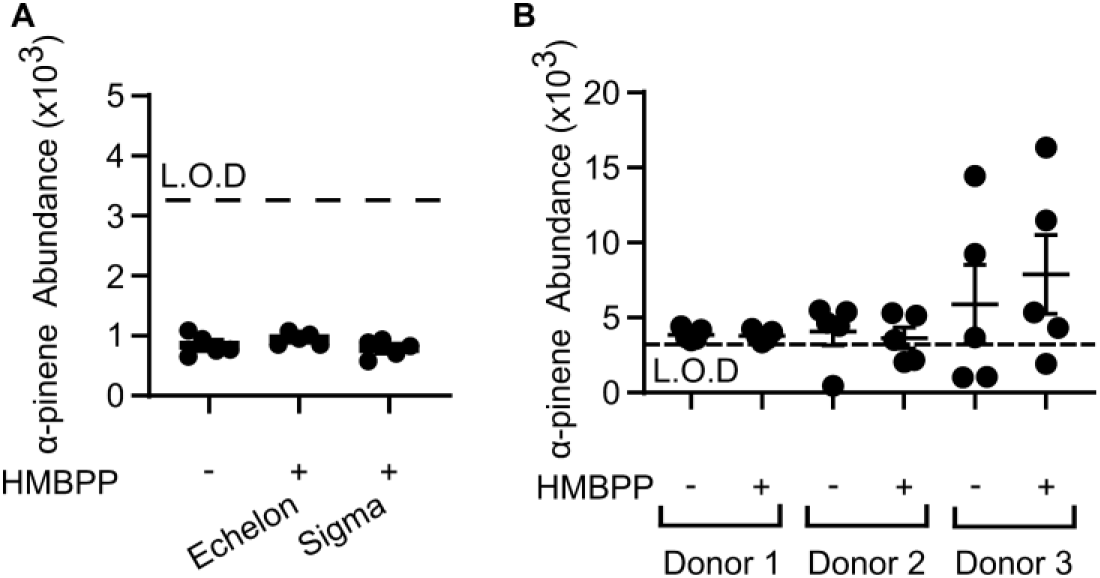
(A) α-pinene abundance in HMBPP from Echelon Biosciences, Sigma Aldrich, or vehicle control (water). (B) Erythrocyte α-pinene abundance following treatment with HMBPP or vehicle control (water). Values are not significantly different by Mann-Whitney U test (Donor 1: p = 0.841, Donor 2: p = 0.548, Donor 3: p = 0.420). All assays performed with n = 5.

In the course of the above experiments, we noted substantial donor-to-donor variability in the endogenous levels of α-pinene present in a given erythrocyte culture. We therefore secured erythrocytes from an additional 3 independent, unrelated donors and quantified α-pinene levels as before. We find that α-pinene levels are strongly dependent on donor identity and range widely among our 6 donors (Fig 3). We find that blood α-pinene concentrations range from 0.37 ng/mL −2.57 ng/mL (mean and standard deviation, 0.91 +/− 0.84 ng/mL). While biosynthesis of α-pinene has not been documented in humans, α-pinene is a volatile component of several common dietary plants, suggesting that one explanation for the variability in α-pinene levels is due to the variability in diet of individual donors. Alternatively, α-pinene may be synthesized by members of the human microbiome that may contribute to endogenous α-pinene levels.

**Figure 3.**
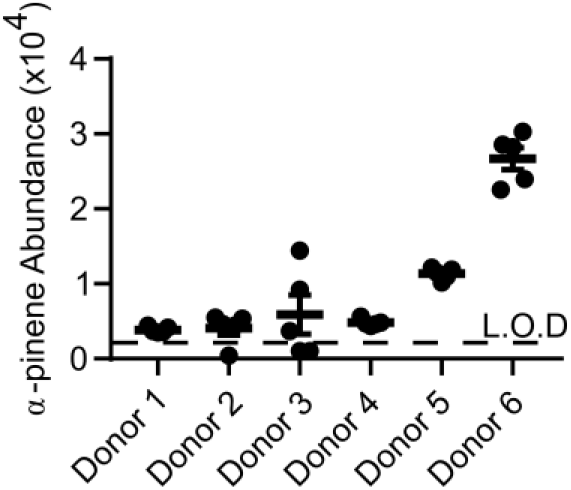
α-pinene abundance in the headspace of untreated human erythrocytes, n = 5.

To reconcile our findings with previous studies that had reported HMBPP-induced α-pinene release, we hypothesized that the previously described variability may be due to a loss of the volatile α-pinene through diffusion, following repeated sampling of the same sample over time. To test this hypothesis, we filled a single air-tight sample tube with erythrocytes from a single donor. At t=0, we removed 1 mL of erythrocytes from the conical tube and measured the headspace concentration of α-pinene according to our previous assay. We left the remainder of the sample sealed (Fig 4A). We repeated this process for a total of 10 runs, allowing the tube of erythrocytes to reequilibrate for one hour between sampling. We find that α-pinene levels decrease by 25-60% as a result of repeated sampling (Fig 4B). As expected given its vapor pressure (4.75 mm Hg at 25 °C), α-pinene is in a vapor-liquid equilibrium^30^. Each time our pooled erythrocytes are uncapped and sampled, vaporous α-pinene diffuses away, and a new vapor-liquid equilibrium is established. The total concentration of liquid α-pinene is thus depleted over time, thereby resulting in a reduced pool of α-pinene in each subsequent sampling. To confirm that this is not unique to α-pinene naturally absorbed within erythrocytes, we supplemented erythrocytes with 10 ng/mL (73.4 nM) α-pinene and found that α-pinene levels drop by 66-80% over the course of repeated sampling (Fig 4C).

**Figure 4.**
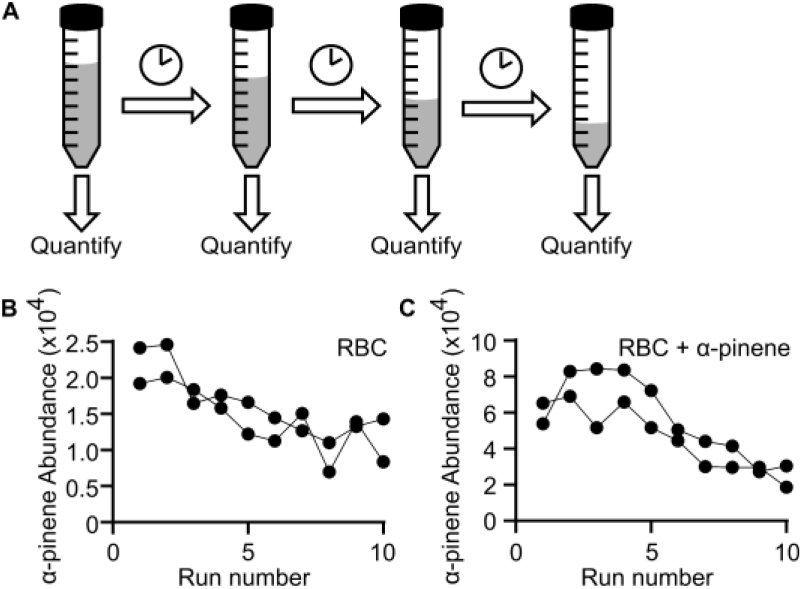
(A) Schematic of repeated sampling mechanism. Time between each sampling is one hour. (B, C) Headspace concentration of α-pinene over untreated human erythrocytes (B) or erythrocytes supplemented with 10 ng/mL commercial α-pinene as a function of GC-MS run.

While run-to-run variability and the run-order effect is a commonly observed problem for mass spectrometry, our results highlight an additional precaution that needs to be taken when sampling biologically generated volatiles. All samples should be aliquoted and sealed in an air-tight container prior to the start of the experiment, as repeated sampling from the same container will result in artificially decreased volatile concentration over time. Investigators should continue to control for run-order effects by randomizing the order in which samples are run.

While mounting evidence suggests that *Plasmodium* infection alters host odor profiles and results in increased mosquito attraction, the mechanism by which this occurs remains unclear. One class of molecules, terpenes, notably α-pinene, has been repeatedly highlighted for being both mosquito attractive and enriched during *Plasmodium* infection. The metabolic origin of terpenes during *Plasmodium* infection remains unclear as mammals do not express orthologs of the terpene synthases required for terpene production. Here, we establish that endogenous levels of α-pinene are present in human erythrocytes. While α-pinene levels from erythrocytes from a single donor sample are highly reproducible, α-pinene levels vary widely by erythrocyte donor. While the source of erythrocyte α-pinene remains enigmatic, it is possible that α-pinene may be dietary in origin, explaining the donor-to-donor variability that we observe.

While HMBPP-mediated α-pinene release has been previously reported^25^, in contrast we do not find evidence that the headspace of HMBPP-treated erythrocytes contains increased levels of α-pinene. HMBPP-treated erythrocytes also appear more mosquito attractive than untreated erythrocytes^25^. Human erythrocytes bind several chemokines^31,32^ and human Vγ9Vδ2-T cells actively respond to HMBPP^28^, raising the possibility that HMBPP exposure of erythrocytes may result in other properties that increase mosquito attraction, independent of α-pinene release. CO_2_ emission from erythrocytes has also been reported to be elevated upon HMBPP exposure. As CO_2_ is also a mosquito semiochemical^33,34^, elevated CO_2_ levels could be responsible for mosquito attraction to HMBPP-treated erythrocytes. However, supplementation of 5 ppm CO_2_ to untreated erythrocytes was not sufficient to sway mosquitoes from HMBPP-treated erythrocytes.

Subsequent experiments are needed to identify the origin of *Plasmodium* infection-associated volatiles. Infection of germ-free animal models may be valuable in discerning volatiles that arise from microbiome vs. host or *Plasmodium* parasite metabolism. Identification of either human or malarial terpene synthases or metabolic labeling studies are required to understand the origin of *Plasmodium* infection-associated terpenes. Carefully controlled dietary recall studies are necessary to understand whether erythrocyte endogenous α-pinene is biosynthesized by humans or human microbiome members.

## Materials and Methods

### Materials and reagents

(E)-4-hydroxy-3-methyl-but-2-enyl pyrophosphate, HMBPP, was purchased from both Sigma Aldrich and Echelon Biosciences Incorporated (Salt Lake City, Utah, USA), resuspended at 4 mM in highly purified water, and stored at −80 °C. Human erythrocytes (types A, B, and O, leukocyte reduced and irradiated) were obtained from the St. Louis Children’s Hospital Blood Bank (St. Louis, Missouri, USA).

### Volatile collection and GC-MS analysis

Erythrocytes were washed 3 times with an equal volume of RPMI-1640 media (Sigma-Aldrich, SKU R4130) supplemented with: 27 mM sodium bicarbonate, 11 mM glucose, 5 mM HEPES, 1 mM sodium pyruvate, 0.37 mM hypoxanthine, 0.01 mM thymidine, 10 μg/mL gentamycin, and 0.5% albumax (Thermo Fisher Scientific, 11020039) and stored at 50% hematocrit at 4 °C. When testing responses of erythrocytes to HMBPP and water, erythrocytes were stored as 1.4 mL aliquots in individual 1.5 mL microfuge tubes (Thermo Fisher Scientific, 05-408-129), wrapped in parafilm (Sigma-Aldrick, SKU P7793) and stored at 4 °C. Prior to sampling, 1 mL 50% erythrocytes were transferred to 4 mL glass autosampler tubes (Thermo Fisher Scientific, 03-391-19), closed with screw caps with septa (Thermo Fisher Scientific 03-391-21), and equilibrated at 38 °C for 15 minutes. Following equilibration, 2.5 μL of 4 mM HMBPP (final concentration 10 μM) or purified water were added to the erythrocytes, caps were closed, and parafilm was used to seal the vial. Volatiles were immediately collected from the headspace using solid phase micro-extraction (n=5, randomized order for each sample). Directly before sampling, the Divinylbenzene/Caboxen/Polydimethylsiloxane fiber (Sigma-Aldrich, SKU 57348) fiber was conditioned for 30 minutes at 225 °C in the inlet of an Agilent 7890A gas chromatographer. Headspace sampling occurred over 30 minutes with temperatures maintained at 38 °C. The volatiles were then desorbed onto the injector of the Agilent 7890A gas chromatographer with an Agilent HP-5MS column (30m, 0.25-mm inner diameter, 0.25-μm film thickness) and interfaced with an Agilent 5975C mass spectrometer. The oven program followed a linear temperature gradient, with an initial temperature of 60 °C (held for 2 minutes), a ramp of 10 °C/min until 225 °C, and a final hold for 5 minutes at 225 °C. Helium was used as the carrier gas with a constant flow of 1 mL/min (25.6 cm/sec). The ion source temperature, electron energy, and emission current were set at 230 °C, 70 eV, and 300 μA respectively. α-pinene was identified based on retention time of an analytical standard (Sigma-Aldrich, SKU 80605), and abundance was quantified using the area under the curve of extracted ion 93. Integration was performed in Agilent MassHunter (Version B.05.00 Build 5.0.519.0) using the Agile integrator. To measure the background contamination of HMBPP with α-pinene 2.5 μL of 4 mM HMBPP (final concentration 10 μM) was added to 1 mL of erythrocyte storage media. α-pinene standard curve was generated through the addition of 2.5 μL of commercial α-pinene diluted in hexanes to autosampler tubes containing 1 mL pure water and sampled as with erythrocyte treatments. The limit of detection was defined as 3 times the area under the curve (ion 93) at the retention time for commercial α-pinene in negative controls containing only water and sampled as with erythrocytes.

### Measuring α-pinene time-dependent concentration

To measure α-pinene loss over time, 14 mL of washed erythrocytes were placed in a 15 mL conical (Sigma-Aldrich, SKU CLS430791) and stored at 4°C. For some experiments, analytical α-pinene, diluted in water, was added to a final concentration of 10 ng/mL in erythrocytes at the time of aliquoting. During the experiment, erythrocytes were stored, capped, on ice and iteratively sampled from. Volatile collection and GC-MS analysis proceeded as above.

## References

1. Alavi, Y. et al. The dynamics of interactions between *Plasmodium* and the mosquito: a study of the infectivity of *Plasmodium berghei* and *Plasmodium gallinaceum*, and their transmission by *Anopheles stephensi, Anopheles gambiae* and *Aedes aegypt*. Int. J. Parasitol. 33, 933–43 (2003).

2. Koella, J. C., Sørensen, F. L. & Anderson, R. A. The malaria parasite, *Plasmodium falciparum*, increases the frequency of multiple feeding of its mosquito vector, *Anopheles gambiae*. Proc. Biol. Sci. 265, 763–8 (1998).

3. Lacroix, R., Mukabana, W. R., Gouagna, L. C. & Koella, J. C. Malaria infection increases attractiveness of humans to mosquitoes. PLoS Biol. 3, 1590–1593 (2005).

4. Busula, A. O. et al. Gametocytemia and attractiveness of *Plasmodium falciparum*– infected Kenyan children to *Anopheles gambiae* mosquitoes. J. Infect. Dis. 216, 291–295 (2017).

5. Batista, E. P., Costa, E. F. & Silva, A. A. *Anopheles darlingi (Diptera: Culicidae)* displays increased attractiveness to infected individuals with *Plasmodium vivax* gametocytes. Parasit. Vectors 7, 251 (2014).

6. De Moraes, C. M. et al. Malaria-induced changes in host odors enhance mosquito attraction. Proc. Natl. Acad. Sci. U. S. A. 111, 11079–84 (2014).

7. Cornet, S., Nicot, A., Rivero, A. & Gandon, S. Malaria infection increases bird attractiveness to uninfected mosquitoes. Ecol. Lett. 16, 323–329 (2013).

8. Schaber, C. L. et al. Breathprinting reveals malaria-associated biomarkers and mosquito attractants. J. Infect. Dis. 217, 1553–1560 (2018).

9. Berna, A. Z. et al. Diurnal variation in expired breath volatiles in malaria-infected and healthy volunteers. J. Breath Res. 12, 46014 (2018).

10. De Moraes, C. M. et al. Volatile biomarkers of symptomatic and asymptomatic malaria infection in humans. Proc. Natl. Acad. Sci. 115, 5780–5785 (2018).

11. Berna, A. Z. et al. Analysis of breath specimens for biomarkers of Plasmodium falciparum infection. J. Infect. Dis. 212, 1120–1128 (2015).

12. Robinson, A. et al. Plasmodium-associated changes in human odor attract mosquitoes. Proc. Natl. Acad. Sci. U. S. A. 115, E4209–E4218 (2018).

13. de Boer, J. G. et al. Odours of *Plasmodium falciparum*-infected participants influence mosquito-host interactions. Sci. Rep. 7, 9283 (2017).

14. Kelly, M. et al. Malaria parasites produce volatile mosquito attractants. mBio 6, 1–6 (2015).

15. Stotzky, G. & Schenck, S. Volatile organic compounds and microorganisms. CRC Crit. Rev. Microbiol. 4, 333–82 (1976).

16. Cane, D. E. & Ikeda, H. Exploration and mining of the bacterial terpenome. Acc. Chem. Res. 45, 463–72 (2012).

17. Yamada, Y. et al. Terpene synthases are widely distributed in bacteria. Proc. Natl. Acad. Sci. 112, 857–862 (2015).

18. Ditengou, F. A. et al. Volatile signalling by sesquiterpenes from ectomycorrhizal fungi reprogrammes root architecture. Nat. Commun. 6, 6279 (2015).

19. Mann, F. M. & Peters, R. J. Isotuberculosinol: the unusual case of an immunomodulatory diterpenoid from Mycobacterium tuberculosis. Medchemcomm 3, 899–904 (2012).

20. Beran, F. et al. Novel family of terpene synthases evolved from trans-isoprenyl diphosphate synthases in a flea beetle. Proc. Natl. Acad. Sci. 113, 2922–2927 (2016).

21. Gilg, A. B., Tittiger, C. & Blomquist, G. J. Unique animal prenyltransferase with monoterpene synthase activity. Naturwissenschaften 96, 731–735 (2009).

22. Lancaster, J. et al. De novo formation of an aggregation pheromone precursor by an isoprenyl diphosphate synthase-related terpene synthase in the harlequin bug. Proc. Natl. Acad. Sci. 115, E8634–E8641 (2018).

23. Wondwosen, B. et al. Sweet attraction: sugarcane pollen-associated volatiles attract gravid Anopheles arabiensis. Malar. J. 17, 90 (2018).

24. Wondwosen, B. et al. Rice volatiles lure gravid malaria mosquitoes, *Anopheles arabiensis*. Sci. Rep. 6, 37930 (2016).

25. Emami, S. N. et al. A key malaria metabolite modulates vector blood seeking, feeding, and susceptibility to infection. 4563, 1–9 (2017).

26. Lange, B. M., Rujan, T., Martin, W. & Croteau, R. Isoprenoid biosynthesis: the evolution of two ancient and distinct pathways across genomes. Proc. Natl. Acad. Sci. U. S. A. 97, 13172–7 (2000).

27. Goldstein, J. L. & Brown, M. S. Regulation of the mevalonate pathway. Nature 343, 425–30 (1990).

28. Eberl, M. et al. Microbial isoprenoid biosynthesis and human γδ T cell activation. FEBS Lett. 544, 4–10 (2003).

29. Rigau, M. et al. Butyrophilin 2A1 is essential for phosphoantigen reactivity by γδ T cells. Science. 367, eaay5516 (2020).

30. Daubert, T. & Danner, R. Physical and Thermodynamic Properties of Pure Chemicals: Data Compilation. (Washington, DC: Taylor & Francis, 1989).

31. Darbonne, W. C. et al. Red blood cells are a sink for interleukin 8, a leukocyte chemotaxin. J. Clin. Invest. 88, 1362–9 (1991).

32. Neote, K., Darbonne, W., Ogez, J., Horuk, R. & Schall, T. J. Identification of a promiscuous inflammatory peptide receptor on the surface of red blood cells. J. Biol. Chem. 268, 12247–9 (1993).

33. Omondi, B. A., Majeed, S. & Ignell, R. Functional development of carbon dioxide detection in the maxillary palp of *Anopheles gambiae*. J. Exp. Biol. 218, 2482–2488 (2015).

34. Dekker, T. Carbon dioxide instantly sensitizes female yellow fever mosquitoes to human skin odours. J. Exp. Biol. 208, 2963–2972 (2005).

